# Green Jackfruit Flour Prevents Metabolic Dysfunction-Associated Steatohepatitis and Progression to Hepatocellular Carcinoma via the AMPK and MAPK Signaling Pathways

**DOI:** 10.1101/2024.05.23.595466

**Authors:** Diwakar Suresh, Bharathwaaj Gunaseelan, Akshatha N. Srinivas, Amith Bharadwaj, James Joseph, Megha, Thomas Varghese, Suchitha Satish, Deepak Suvarna, Prasanna K. Santhekadur, Saravana Babu Chidambaram, Ajay Duseja, Divya P. Kumar

## Abstract

Metabolic dysfunction-associated steatotic liver disease (MASLD), encompassing metabolic-dysfunction associated steatotic liver (MAFL) and steatohepatitis (MASH), which further progresses to hepatocellular carcinoma (HCC), is a serious public health concern. Given the paucity of approved therapeutic strategies for this lifestyle disorder, dietary interventions may prove effective. We evaluated how green jackfruit flour (JF) prevents MASH and progression to HCC and its underlying mechanisms. The study utilized two murine models that mimicked human MASLD disease: (i) a diet-induced MASH model; (ii) a MASH-HCC model induced by diet and a very low dose of CCl_4_. C57Bl/6 mice were fed with chow (CD) or western diet (WD) with normal (NW) or sugar water (SW) for 12 weeks, then randomized to receive either 5 kcal% green jackfruit flour (JF) or an equal volume of placebo flour (PB). The biochemical, histological, and molecular analyses were assessed. JF significantly reduced body weight, liver injury, insulin resistance, and alleviated obesity, steatosis, inflammation, fibrosis, and tumor development in WDSW or WDSW/CCl_4_ mice compared to placebo groups. Furthermore, JF activated AMPK (AMP-activated protein kinase) and inhibited MAPK (mitogen-activated protein kinase) signaling pathways in MASH and MASH-HCC experimental models, respectively. This was supported by sodium propionate treatment, the primary short-chain fatty acid entering the liver from JF’s soluble fiber microbial fermentation, which also regulated AMPK and MAPK signaling in cellular models of MASH and HCC, respectively. Hence, our findings present strong evidence of JF’s therapeutic potential in the prevention of MASH and MASH-HCC, warranting further investigation of JF’s efficacy as a dietary intervention in clinical trials.

## 1. Introduction

Metabolic Dysfunction-Associated Steatotic Liver Disease (MASLD), formerly known as Nonalcoholic Fatty Liver Disease (NAFLD), has emerged as a major worldwide health concern inextricably linked to metabolic abnormalities such as obesity, insulin resistance, and dyslipidemia [1,2]. The MASLD spectrum encompasses a wide array of hepatic manifestations, spanning from benign steatosis (MASL, formerly known as non-alcoholic fatty liver, NAFL) to the progressive inflammatory condition, steatohepatitis (MASH, formerly known as non-alcoholic steatohepatitis, NASH), which can further advance to fibrosis, cirrhosis, and ultimately, liver malignancies [2]. Approximately 30% of the global adult population grapples with MASLD, with projections suggesting a doubling in prevalence within the next decade [3,4]. A notable percentage of affected individuals progress to MASH and associated hepatic tumors, contributing to the escalating incidence of non-viral hepatocellular carcinoma (HCC) [5]. Given the patient heterogeneity and multifarious extrahepatic contributing factors, diagnosing and treating MASLD poses significant challenges, underscoring the importance of adopting a holistic approach [6]. Very recently, in March 2024, Resmetirom (Rezdiffra, Madrigal Pharmaceuticals) was approved by the US Food and Drug Administration (FDA) as the first drug for the treatment of metabolic dysfunction-associated steatohepatitis (MASH), in addition to diet and exercise, and is currently not available worldwide [7]. Hence, diet and physical activity are considered the principal components of lifestyle intervention. Regular physical exercise combined with dietary adjustments has demonstrated efficacy in improving liver function [8]. Additionally, adherence to a mediterranean-style diet, consumption of antioxidant-rich foods, as well as the inclusion of black coffee in dietary regimens, have shown promise in ameliorating MASLD symptoms by promoting weight loss, improving insulin sensitivity, reducing inflammation, and regulating glucose and lipid metabolism [9,10].

In recent years, there has been growing interest in natural products and dietary interventions for their potential therapeutic effects in mitigating the progression of MASLD and reducing the risk of HCC [11,12]. *Artocarpus heterophyllus*, commonly known as jackfruit, has garnered increasing attention due to its diverse therapeutic properties across various health conditions [13]. Native to South and Southeast Asia, *A. heterophyllus* belongs to the Moraceae family and has been traditionally utilized for its nutritional value and medicinal benefits. With a rich composition of phytochemicals such as flavonoids, phenolic compounds, and saponins, *A. heterophyllus* exhibits a wide range of biological activities, including antioxidant, anti-inflammatory, antimicrobial, and anticancer properties [14–17]. One of the most intriguing aspects of *A. heterophyllus* is its custom of consuming green unripe jackfruit as a meal, an efficient alternative to rice for regulating blood sugar levels, owing to its soluble fiber, pectin content, and lower glycemic load [18]. Many studies have shown that bioactive components present in both unripe and ripened jackfruit help in mitigating diabetes and obesity-associated metabolic dysfunctions [16,19]. Previous studies by Rao AG et al. show the beneficial impact of incorporating green jackfruit-based flour into the diet as a medical nutrition therapy for type 2 diabetes mellitus, wherein it replaces an equal volume of rice or wheat in daily meals [20]. Similarly, findings from a study by Varughese et al. have shown that a diet with green jackfruit flour can mitigate chemotherapy-induced leukopenia (CIL) in patients undergoing chemotherapy. Notably, the inclusion of green jackfruit flour in the daily diet has been found to prevent CIL in these patients, suggesting its potential as a supportive therapy during cancer treatment [21]. The therapeutic potential of *Artocarpus heterophyllus*, particularly its role in managing metabolic disorders and supporting cancer patients undergoing chemotherapy, underscores its value as a natural remedy with diverse health benefits. However, the efficacy of green jackfruit flour in preventing obesity and MASH remains unexplored. Gaining molecular insights into the therapeutic potential of green jackfruit flour holds promise to enhance its effectiveness in clinical practice.

Given these considerations, this study sets out to explore how green jackfruit flour could potentially prevent obesity, MASH, and progression to HCC, focusing primarily on unraveling the underlying molecular mechanisms. To accomplish this, we have employed both cellular and preclinical murine models of MASH and MASH-HCC. Here we detail the efficacy of 5 kcal% of green jackfruit flour versus placebo flour (wheat flour) in reducing body weight, liver injury, inflammation, fibrosis, and tumor development in experimental mouse models. Our data indicate that the underlying mechanism would be the action of propionate, which is produced by the fermentation of soluble dietary fibers from green jackfruit flour. Propionate modulates the AMPK and MAPK signaling pathways and provides significant protection to the liver against MASH and HCC. Taken together, these findings offer mechanistic insights into the efficacy of green jackfruit flour as a dietary intervention for the prevention of MASH and HCC.

## 2. Materials and Methods

### 2.1 Materials

Green jackfruit flour (Jackfruit 365^TM^) and wheat flour were purchased from Amazon and shipped to Research Diet Inc. (NJ, USA) for the preparation of customized diet. The chow diet and western diet were prepared with 5 kcal% of green jackfruit flour or wheat flour (placebo flour). The Accu-Chek Glucometer was procured from Roche Diagnostics (Rz, Switzerland). AST, ALT, triglycerides, and total cholesterol kits were from Monlab tests (Spain); oil red o stain, picrosirius red stain, sodium propionate, free fatty acids, TRIzol, and RIPA buffer were from Sigma-Aldrich (St. Louis, USA). Verso cDNA synthesis kit and DyNamo Colorflash SYBR green kit from Thermo Fisher Scientific (USA). All the cell culture chemicals were from Invitrogen (USA) or HiMedia Laboratories (India). Western blotting materials were from Bio-Rad and western bright ECL HRP substrate from Advansta (USA). Primers were procured from IDT (Iowa, USA), and primary antibodies to SREBP1, pErk1/2, Erk1/2, pJNK, and JNK were from Santa Cruz Biotechnology, Inc. (CA, USA); antibodies to pP38 MAPK, P38 MAPK, ACC, FASN, CK19, F4/80, and β-actin were from Cell Signaling Technologies (USA).

### 2.2 Experimental model

C57Bl/6 mice (Charles River Laboratories) aged 5–6 weeks were procured from Hylasco Biotechnology Pvt. Ltd. These mice were housed in compliance with previously established protocols [22] within the animal facility managed by the Centre for Experimental Pharmacology and Toxicology, JSS AHER. The mice were maintained under controlled environmental conditions, including a 12-hour light-dark cycle and temperatures ranging between 21 and 24 °C. The parameters such as body weight, food intake, and water consumption were regularly monitored throughout the study period. All procedures involving animals were conducted following approved protocols and adhering to the guidelines and regulations outlined by the Committee for Control and Supervision of Experiments on Animals (CPCSEA), Government of India (JSSAHER/CPT/IAEC/165/2023).

### 2.3 Study design and dietary intervention

Following acclimatization to the housing conditions, the mice were randomly divided into different dietary interventions for both MASH and MASH-HCC models.

#### MASH model

The mice were fed with either a chow diet with normal drinking water (CDNW) or a western diet comprising high fat, high sucrose, and high cholesterol (21% fat, 41% sucrose, and 1.25% cholesterol by weight) along with sugar water (WDSW; 18.9 g/L d-glucose and 23.1 g/L d-fructose) ad libitum for 12 weeks. After 12 weeks, mice were fed with either 5 kcal% of green jackfruit flour or an equal volume of placebo flour diet for an additional 12 weeks. Mice were divided into the following groups: (i) CDNW+ placebo flour; (ii) CDNW+ green jackfruit flour; (iii) WDSW+ placebo flour; and (iv) WDSW+ green jackfruit flour. At the end of the dietary intervention, mice were sacrificed, and blood and tissues were collected for further analysis (**Figure 1A**).

**Figure 1:**
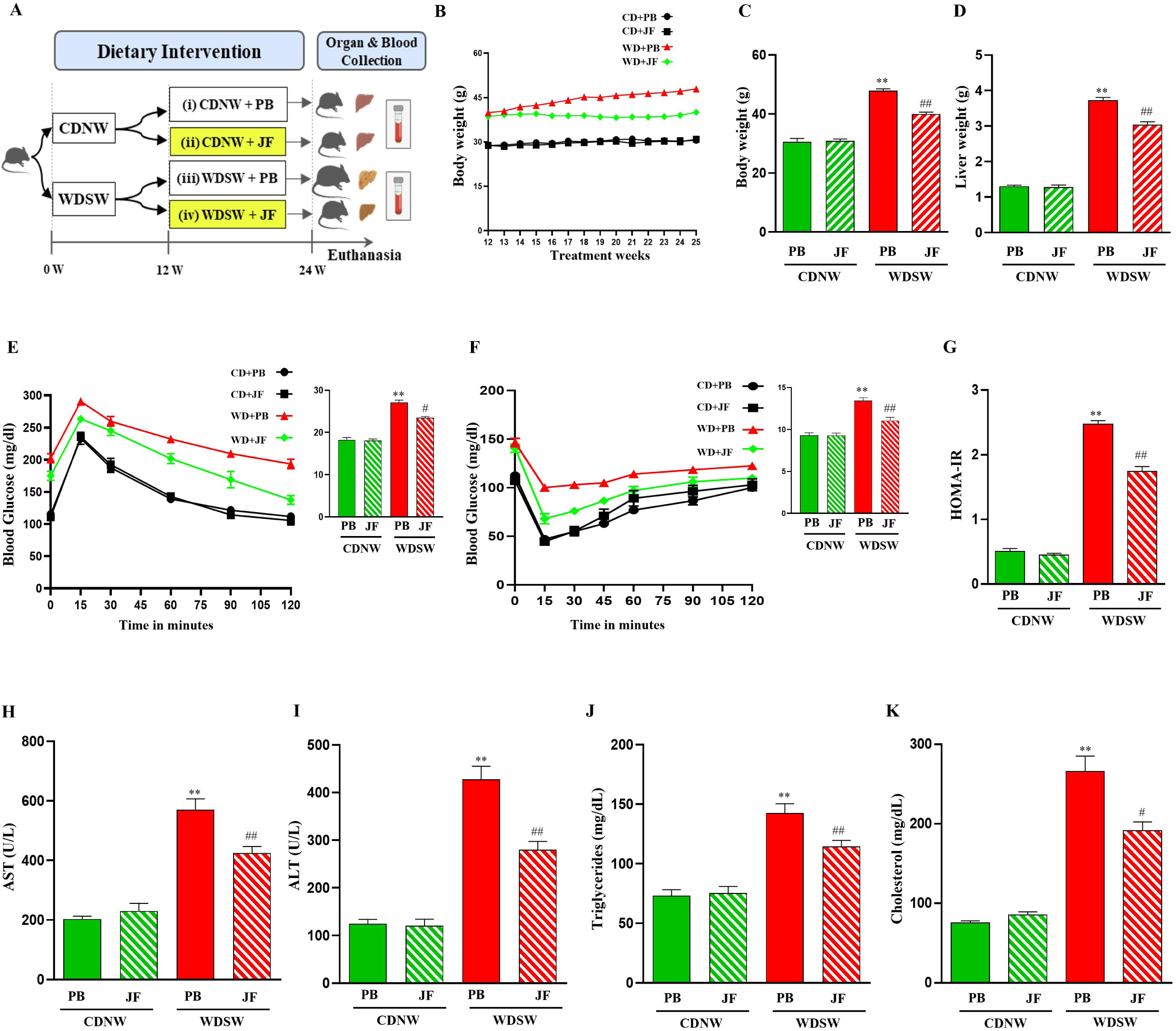
Green jackfruit flour prevents obesity, insulin resistance and liver injury in the experimental MASH model. Mice were fed with either CDNW or WDSW for 12 weeks. Further, mice were randomized into four groups based on dietary interventions. Mice were fed with CDNW with placebo flour (PB) or green jackfruit flour (JF) and WDSW with placebo flour or green jackfruit flour for an additional 12 weeks (A). Body weight during 12 weeks of dietary intervention (B), body weight at the end of the treatment (C), and liver weight (D) were measured. GTT (E) and ITT (F) were performed on overnight and 4-5 hours-fasted mice, respectively. The bar graphs represent the area under the curve (AUC) for different groups. HOMA-IR was calculated using the values of fasting glucose and fasting insulin (G). Serum levels of the liver enzymes AST (H), ALT (I), triglycerides (J), and cholesterol (K) were measured. All the data are expressed as mean±SEM for 6-8 mice per group. **p<0.001 compared to CDNW PB; ^##^p<0.001 or ^#^p<0.05 compared to WDSW PB. CDNW, chow diet normal water; WDSW, western diet sugar water PB, placebo flour; JF, green jackfruit flour; GTT, glucose tolerance test; ITT, insulin tolerance test; HOMA-IR, homeostatic model assessment for insulin resistance; AST, aspartate aminotransferase; ALT, alanine aminotransferase.

#### MASH-HCC model

One group of mice received a standard chow diet along with normal water (CDNW). Another group was subjected to a western diet characterized by high levels of fat, sucrose, and cholesterol, comprising 21% fat, 41% sucrose, and 1.25% cholesterol by weight, along with sugar water (WDSW) containing 18.9 g/L d-glucose and 23.1 g/L d-fructose. To accelerate the HCC progression, all mice were treated with a very low dose of CCl_4_ (0.2 μl/g) of body weight dissolved in corn oil throughout the study, as described previously [23]. After 12 weeks, mice were fed with either 5 kcal% of green jackfruit flour or an equal volume of placebo flour diet for an additional 12 weeks, divided into 4 groups: (i) CDNW/CCl_4_+ placebo flour; (ii) CDNW/ CCl_4_+ green jackfruit flour; (iii) WDSW/CCl_4_+ placebo flour; and (iv) WDSW/CCl_4_+ green jackfruit flour. At the end of 24 weeks, mice were euthanized for blood and tissue collection (**Figure 3A**).

**Figure 2:**
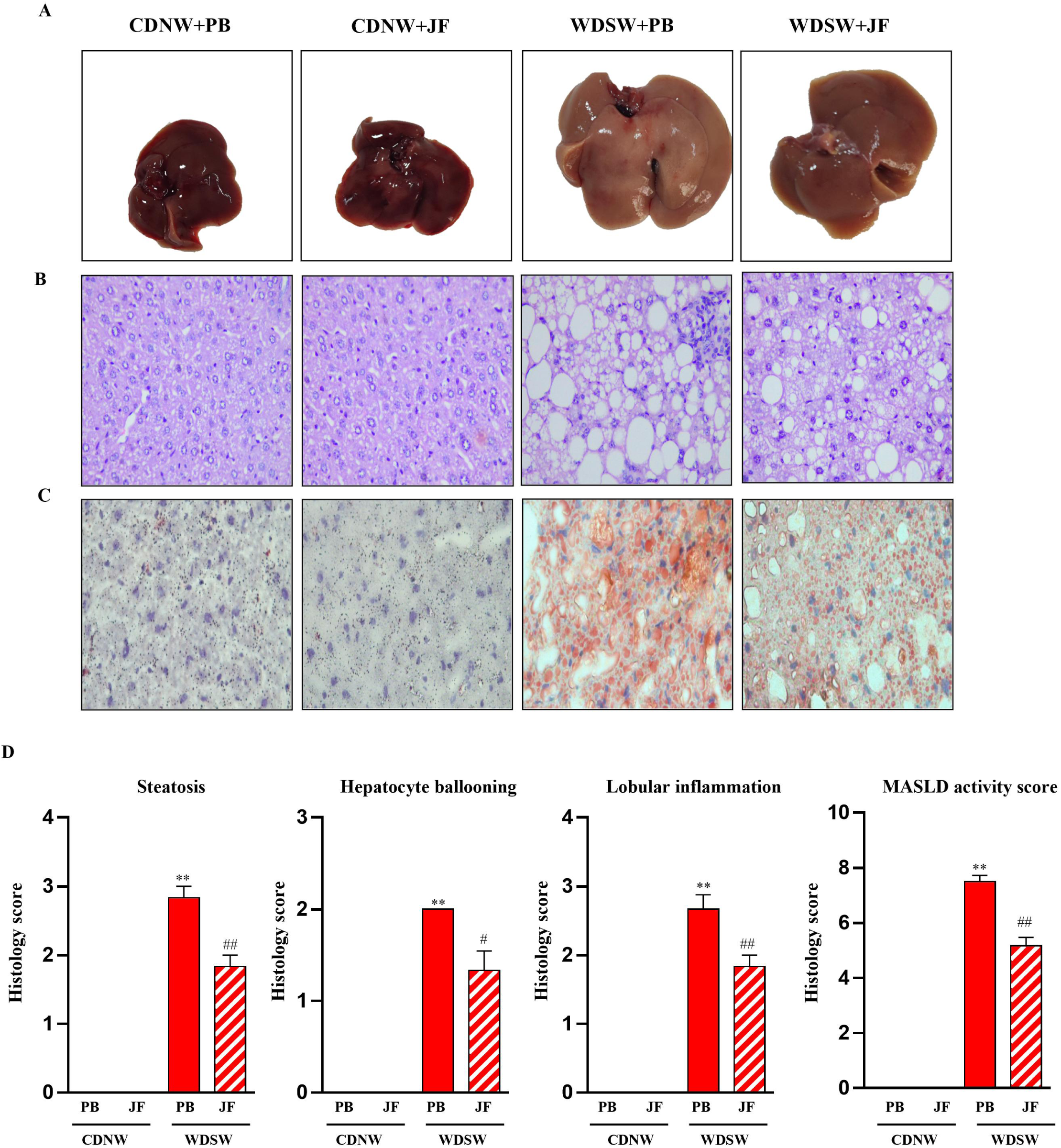
Histological features of mice upon green jackfruit flour dietary intervention in the experimental MASH model. Representative macroscopic and microscopic images of liver from CDNW and WDSW fed with either placebo flour (PB) or green jackfruit flour (JF). Liver picture (A), hematoxylin & eosin staining (400X) (B), Oil red O staining depicting intracellular lipid accumulation (C), histology score for (D) steatosis, (E) hepatocyte ballooning, (F) lobular inflammation, and (G) MASLD activity score were quantified. All the data are expressed as mean±SEM for 6-8 mice per group. **p<0.001 compared to CDNW PB; ^##^p<0.001 or ^#^p<0.05 compared to WDSW PB. CDNW, chow diet normal water; WDSW, western diet sugar water; PB, placebo flour; JF, green jackfruit flour; MASLD, metabolic dysfunction-associated steatotic liver disease.

**Figure 3:**
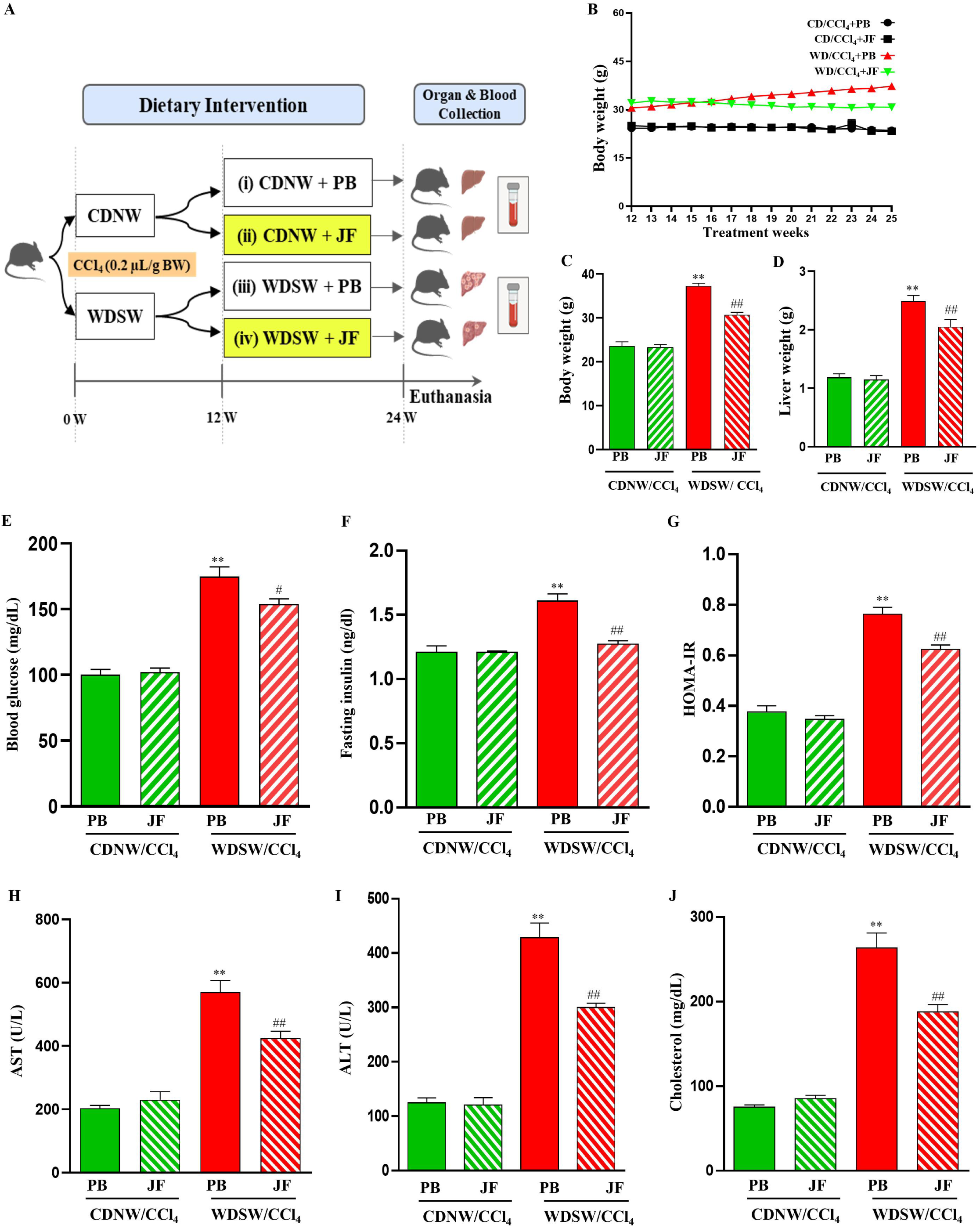
Green jackfruit flour prevented insulin resistance and liver injury in the experimental MASH-HCC model. Mice were fed with either CDNW or WDSW for 12 weeks, along with a very low dose of CCl_4_ given intraperitonially every week. Further, mice were divided into four groups based on dietary interventions. Mice were fed with CDNW with placebo flour (PB) or green jackfruit flour (JF) and WDSW with placebo flour or green jackfruit flour for an additional 12 weeks with no changes in CCl_4_ treatment (A). Body weight during 12 weeks of dietary intervention (B), body weight at the end of the treatment (C), and liver weight (D) were measured. Fasting blood glucose (E), fasting insulin (F), and HOMA-IR (G) were estimated. Serum levels of the liver enzymes AST (H), ALT (I), and cholesterol (J) were measured. All the data are expressed as mean±SEM for 6-8 mice per group. **p<0.001 compared to CDNW/CCl_4_ PB; ^##^p<0.001 or ^#^p<0.05 compared to WDSW/CCl_4_ PB. CDNW, chow diet normal water; WDSW, western diet sugar water; PB, placebo flour; JF, green jackfruit flour; HOMA-IR, homeostatic model assessment for insulin resistance; AST, aspartate aminotransferase; ALT, alanine aminotransferase.

### 2.4 Biochemical analysis

Blood samples were obtained via retro-orbital puncture from mice that had undergone an overnight fast before euthanasia. Subsequently, the samples were allowed to clot at room temperature and were centrifuged at 1500 g for 15 minutes at 4°C. The serum was collected and stored at −80°C until analysis. Serum levels of AST, ALT, cholesterol, and triglycerides were quantified using kits following the manufacturer’s guidelines.

### 2.5 Glucose Tolerance and Insulin Tolerance Tests

An overnight fasting glucose tolerance test (GTT) was conducted on mice by initially measuring baseline blood glucose levels using an Accu-Check Glucometer. Glucose (1 g/kg body weight) dissolved in phosphate-buffered saline was then intraperitoneally administered to the mice. Following glucose injection, blood glucose levels were monitored at various time points (0, 15, 30, 60, 90, and 120 minutes). Similarly, an insulin tolerance test (ITT) was performed after fasting the mice for 5–6 hours and measuring baseline glucose levels similar to the GTT. Mice were intraperitoneally injected with insulin (0.75 U/kg body weight), and blood glucose levels were measured at different intervals (0, 15, 30, 45, 60, 90, and 120 minutes) thereafter. Both GTT and ITT were conducted before the dietary intervention (12 weeks) and again at the end of the study (24 weeks).

### 2.6 ELISA

The serum levels of fasting insulin mice were determined using the ELISA kits (Krishgen Biosystems) according to the manufacturer’s protocol.

### 2.7 Histological analysis

Formalin-fixed and paraffin-embedded mouse liver tissues were stained for hematoxylin and eosin to evaluate the tissue morphology. Tissues were scored blindfolded for the degree of steatosis, hepatocyte ballooning, and lobular inflammation by an expert pathologist at JSS Hospital. MASLD activity scoring was calculated using these scores according to the FLIP algorithm and NASH-clinical research network (CRN) criteria [24,25].

### 2.8 Oil Red O staining

Frozen liver tissues were used for the Oil red O staining. 12-μm-thick cryosections were fixed with 10% formaldehyde for 5 minutes and washed with distilled water. The slides were incubated with 60% isopropanol, followed by the Oil red O solution for one hour. The excess stain was rinsed off with distilled water, and the nucleus was stained with hematoxylin. Slides were mounted, and images were taken with a Leica microscope.

### 2.9 Sirius Red staining

The formalin-fixed, paraffin-embedded liver tissue sections were deparaffinized and rehydrated using xylene, followed by ethanol. Further, the slides were stained with hematoxylin for 10 minutes. After water wash, the slides were subjected to sirius red solution staining for 1 hour at room temperature. The excess stain was rinsed off by washing the slides in 0.5% acidified water for 5 minutes. The slides were mounted, and images were captured using a Leica microscope.

### 2.10 Immunohistochemistry

Following deparaffinization and rehydration, the formalin-fixed, paraffin-embedded liver tissues were incubated in a citrate buffer of pH 6 at 94^0^C for 15 minutes for antigen retrieval. The tissue sections were incubated with 3% hydrogen peroxide for 5 minutes and blocked with normal goat serum for 2 hours. The primary antibody (F4/80-1:100) was added to the liver tissue section and incubated overnight at 4^0^C in a humidified chamber. Polyexcel HRP/DAB detection system one step (PathnSitu, Biotechnologies) was used for developing the signals and hematoxylin for counterstaining the nucleus. All the images were taken using a Leica microscope.

### 2.11 Cell culture

Human hepatic cells, HepG2 cells (procured from the American Type Culture Collection, ATCC) were cultured in DMEM containing 1 g/L glucose supplemented with 10% fetal bovine serum. Human HCC cells, Hep3B (procured from the American Type Culture Collection, ATCC), and QGY-7703 cells (a kind donation from Dr. Devanand Sarkar, Virginia Commonwealth University, USA), were cultured in DMEM containing 4.5 g/L glucose supplemented with 10% fetal bovine serum. Cells were grown in respective culture media containing L-glutamine and 100 U/ml penicillin-streptomycin and incubated at 37^0^C in 5% CO_2_.

#### *In vitro* model for steatosis

HepG2 cells were cultured up to 50% confluency in cell culture plates. The cells were treated with 0.5 mM sodium oleate (SO) or 0.5 mM sodium palmitate (SP) dissolved in methanol in 5% fatty acid-free BSA containing culture media. After 24 hours of treatment, the cells were assessed for lipid droplets using Oil red O staining.

### 2.12 Cell viability assay

The cell viability was determined by the WST-1 assay as per manufacturer’s instructions. Different doses (0, 0.5, 1, 2, and 4 mM) of sodium propionate (NaP) were treated to HepG2 cells or Hep3B cells for 24 hrs and 48 hrs and cell viability was assessed. The data is represented as percent cell viability after normalizing with control (no treatment).

### 2.13 RNA Isolation and Quantitative Real-time PCR

Total RNA was extracted from frozen liver tissue and cell pellets using TRIzol following the manufacturer’s instructions. The concentration and quality of the RNA were assessed using a NanoDrop spectrophotometer. Further, the RNA was reverse transcribed into complementary DNA (cDNA) using the Verso cDNA synthesis kit following the manufacturer’s instructions. Quantitative real-time PCR was performed utilizing a Rotor-Gene Q 5plex HRM system (QIAGEN, Hilden, Germany). The reactions were conducted using a DyNamo Colorflash SYBR green kit with 0.5 mM primers (IDT) and 50 ng of cDNA in a 20 μl reaction volume. The relative fold change in mRNA levels was calculated as 2^-ΔΔCt^ and normalized to the endogenous control, β-actin. The validated primer sequences used in the study are given in **Supplementary Table 1.**

### 2.14 Western blotting

Fresh lysates were prepared from both cells and liver tissues using RIPA buffer containing protease and phosphatase inhibitors. After centrifugation of the homogenate at 13,000 rpm for 10 minutes at 4°C, protein supernatants were collected. The protein concentration of these lysates was determined using Bradford’s protein estimation method. Equal amounts of lysates (30 μg) were loaded onto SDS-PAGE gels for protein separation, followed by transfer onto a nitrocellulose membrane. The membranes were then blocked with 5% non-fat skim milk for 1 hour and subsequently incubated overnight at 4°C with primary antibodies targeting specific proteins (pAMPK, AMPK, ACC, FASN, SREBP1, pErk1/2, Erk1/2, pJNK, JNK, pP38 MAPK, P38 MAPK, and β-actin). After primary antibody incubation, the membrane was treated with a secondary antibody for 2 hours at room temperature. Blots were visualized using Western Bright ECL HRP substrate in the UVitec Alliance Q9 chemiluminescence imaging system. Image analysis was performed using ImageJ software, and bands were normalized with corresponding internal controls.

### 2.15 Data analysis

The results were expressed as the mean ± standard error of the mean (SEM). Statistical significance was assessed using either the student’s t-test for comparisons between two groups or the analysis of variance (ANOVA) with post hoc Bonferroni correction for multiple comparisons. All statistical analyses were conducted using GraphPad Prism software (version 6). Values of p < 0.05 (*or ^#^) or < 0.001 (** or ^##^) were considered statistically significant.

## 3. Results

### 3.1 Green Jackfruit flour prevents obesity, metabolic dysfunction-associated steatosis and steatohepatitis

To assess the effect of green jackfruit flour on metabolic dysfunction-associated steatohepatitis, C57Bl/6 mice were fed with either a chow diet normal water (CDNW) or a western diet sugar water (WDSW) for 12 weeks. Following this, the mice were further randomized into four groups based on dietary interventions wherein we used 5 kcal% of green jackfruit flour or an equal volume of placebo flour: CDNW+ placebo flour (CDNW+PB), CDNW+ green jackfruit flour (CDNW+JF), WDSW+ placebo flour (WDSW+PB), and WDSW+ green jackfruit flour (WDSW+JF) **(Figure 1A)**. Mice receiving WDSW+JF exhibited a significant reduction in body weight compared to WDSW+PB **(Figure 1B and 1C)**. Similarly, the liver weight was also reduced in WDSW+JF mice **(Figure 1D)**. Notably, there was no difference in the calorie intake between the groups. Furthermore, WDSW+JF mice showed reduced fasting glucose and insulin levels **(Supplementary Figure 1A-1D)**. The response to the glucose tolerance test (GTT) and insulin tolerance test (ITT) indicated that the WDSW+PB mice showed persistent hyperglycemia and insulin resistance compared to the CDNW+PB mice. On the contrary, WDSW+JF mice showed reduced glucose levels and improved insulin sensitivity **(Figure 1E-1G)**. Additionally, JF reduced AST, ALT, triglycerides, and cholesterol in WDSW mice compared to placebo **(Figure 1H and 1K)** but had no effect in the CDNW mice.

In line with the improved metabolic parameters, WDSW+JF mice demonstrated improved liver health, which was significantly evident in the H/E stained-liver sections **(Figure 2A and 2B)**. JF also reduced lipid accumulation in WDSW mice compared to placebo **(Figure 2C)**. These findings were corroborated by histological examinations, which indicated severe macrovesicular and microvesicular steatosis (2.83±0.17), hepatocyte ballooning (1.83±0.16), and inflammation (2.67±0.21) in WDSW+PB mice with a considerable MASLD score (7.5±0.22). In contrast, WDSW+JF mice showed a significant reduction in pathological indicators, including macrovesicular steatosis (1.86±0.29), hepatocyte ballooning (1.33±0.21), lobular inflammation (1.85±0.16), and MASLD score (5.16±0.20), indicating a potential improvement in MASH hallmarks **(Figure 2D-2G)**. In summary, these data suggest that green jackfruit flour protects the liver against WDSW-induced liver damage and insulin resistance in MASH.

### 3.2 Protective effects of green jackfruit flour against liver damage in experimental MASH-HCC

In this study, we have employed two different preclinical murine models: (i) MASH model and (ii) MASH-HCC model. We investigated the protective effects of green jackfruit flour against MASH-associated HCC by employing C57Bl/6 mice fed with either chow diet normal water (CDNW) or western diet sugar water (WDSW) supplemented with a very low dose of CCl_4_ intraperitoneally once a week throughout the study. Upon confirming MASH at the 12^th^ week, the mice were further randomized into four groups based on dietary interventions for an additional 12 weeks wherein we used 5 kcal% of green jackfruit flour or an equal volume of placebo flour: CDNW/CCl4+ placebo flour (CD/CCl_4_+PB), CDNW/CCl4+ green jackfruit flour (CD/CCl_4_+JF), WDSW/CCl4+ placebo flour (WD/CCl_4_+PB), and WDSW/CCl4+ green jackfruit flour (WD/CCl_4_+JF) **(Figure 3A)**. As hypothesized, mice receiving WD/CCl_4_+JF exhibited a significant reduction in body weight and liver weight compared to WD/CCl_4_+PB mice **(Figure 3B-3D)**. The food and calorie intake remained consistent across all the study groups (**Supplementary Figure 2).** Similarly, JF prevented liver damage in WD/CCl_4_ mice by reducing fasting blood glucose and insulin levels and HOMA-IR (**Figure 3E-3G)**. There was also significant improvement in the liver function tests (AST, ALT, total cholesterol) in WD/CCl_4_+JF compared to WD/CCl_4_+PB mice **(Figure 3H-3J)**. However, no significant effects were seen in CD/CCl_4_+JF versus CD/CCl_4_+PB mice.

Furthermore, while the WD/CCl_4_+PB mice developed liver tumors (8-10 per mouse) after 24 weeks, there was a significant reduction in tumor development (2-4 per mouse) in the WD/CCl_4_+JF mice **(Figure 4A and 4C)**. The histological analysis of WD/CCl_4_+JF mice also revealed a notable reduction in steatosis (1.83±0.16 vs. 2.67±0.21 in WD/CCl_4_+PB), hepatocyte ballooning (1.16±0.17 vs. 1.83±0.13 in WD/CCl_4_+PB), and lobular inflammation (1.83±0.26 vs. 2.83±0.16 in WD/CCl_4_+PB) with MASLD activity score (4.83±0.16 vs. 7.33±0.21 in WD/CCl_4_+PB) **(Figure 4D-4G)**. Collectively, these findings clearly indicate that green jackfruit flour prevents MASH and the development of tumorigenesis in the liver of WD/CCl_4_ mice compared to placebo flour.

**Figure 4:**
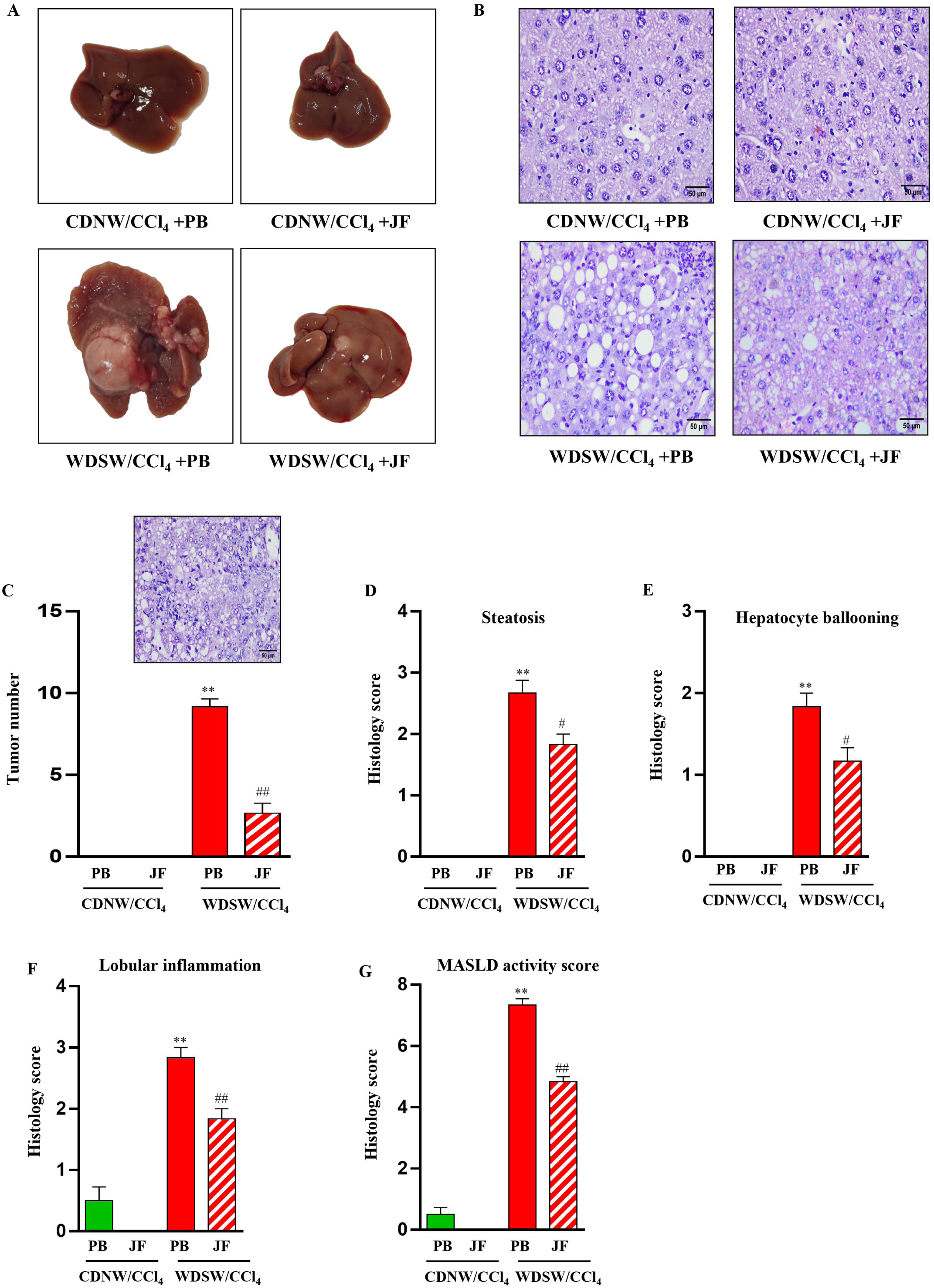
Histological features of mice upon green jackfruit flour dietary intervention in the experimental MASH-HCC model. Representative macroscopic and microscopic images of liver from CDNW/CCl_4_ and WDSW/CCl_4_ fed with either placebo flour (PB) or green jackfruit flour (JF). Liver pictures (A), hematoxylin & eosin staining (400X) (B), tumor number (C), histology score for (D) steatosis, (E) hepatocyte ballooning, (F) lobular inflammation, and (G) MASLD activity score were quantified. All the data are expressed as mean±SEM for 6-8 mice per group. **p<0.001 compared to CDNW/CCl_4_ PB; ^##^p<0.001 or ^#^p<0.05 compared to WDSW/CCl_4_ PB. CDNW, chow diet normal water; WDSW, western diet sugar water; PB, placebo flour; JF, green jackfruit flour; MASLD, metabolic dysfunction-associated steatotic liver disease.

### 3.3 Green Jackfruit flour reduces inflammation, and fibrosis, and prevents hepatocarcinogenesis

The defining characteristics of MASH encompass liver injury and chronic inflammation initiated within the framework of cellular stress, leading to the activation of fibrosis and oncogenic signaling [26]. In this study, we explored the effect of green jackfruit flour on the hallmarks of MASH and MASH-HCC. Proinflammatory cytokines such as TNF-α, IL-1β, and IL-6 were significantly suppressed in WDSW+JF mice compared to the placebo group **(Figure 5A-5C)**. This was further validated through immunostaining for F4/80 to identify the infiltration of monocyte-derived macrophages in the liver. Green jackfruit flour significantly downregulated macrophage infiltration into the liver tissue compared to WDSW+PB mice **(Figure 5D)**. Furthermore, the liver sections were examined for fibrosis by picrosirius red staining, and the WDSW+JF mice exhibited reduced fibrosis **(Figure 5E)**.

**Figure 5:**
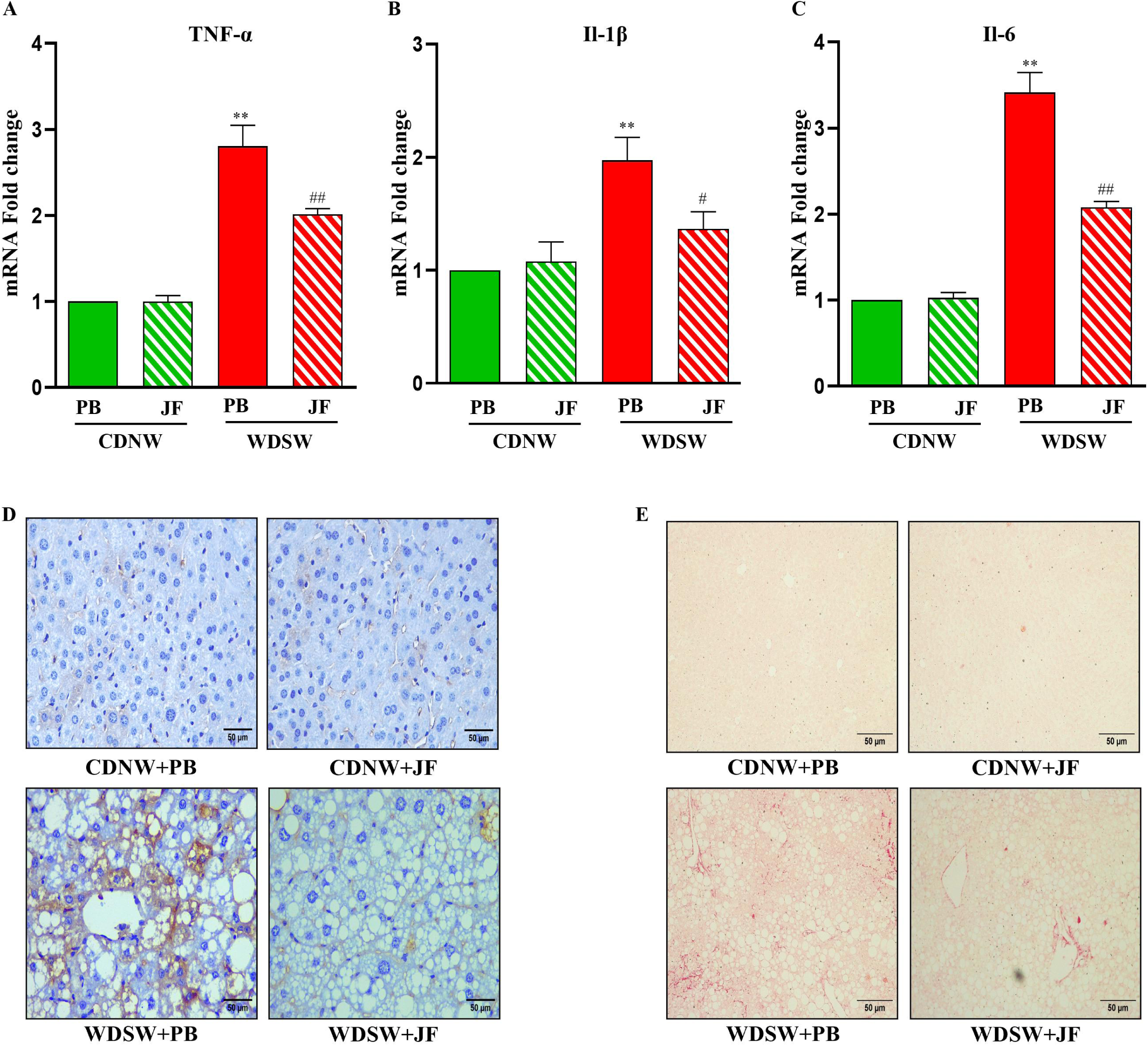
Green jackfruit flour reduces hepatic inflammation and fibrosis in MASH mice. The relative hepatic mRNA expression of (A) TNF-α, (B) IL-1β, and (C) IL-6 was determined by qRT-PCR in CDNW and WDSW mice fed with either placebo flour (PB) or green jackfruit flour (JF). The expression levels were normalized with β-actin. Liver sections were stained for (D) F4/80 antibody and (E) picrosirius red to represent inflammation and fibrosis, respectively. All the data are expressed as mean±SEM for 6-8 mice per group. **p<0.001 or ^#^p<0.05 compared to CDNW PB; ^##^p<0.001 compared to WDSW PB. CDNW, chow diet normal water; WDSW, western diet sugar water; TNF-α, tumor necrosis factor alpha; IL-1β, interleukin 1 beta; IL-6, interleukin 6.

We also examined the proinflammatory cytokines in the experimental MASH-HCC model, and the results were consistent with the MASH model. Green jackfruit flour reduced the expression of pro-inflammatory cytokines such as TNF-α, IL-1β, and IL-6 in mice livers **(Figure 6A-6C).** Similarly, WDSW+JF mice demonstrated a marked reduction in fibrosis compared to WDSW+PB mice **(Figure 6D)**. The hepatic expression of cluster differentiation 31 (CD31) and alpha-fetoprotein (AFP), specific HCC markers was found to be upregulated in WDSW+PB mice. In addition to the reduction in tumor numbers in WDSW+JF mice, we also observed the downregulation of CD31 and AFP in these mice compared to WDSW+PB mice **(Figure 6E and 6F)**. Thus, these results indicate that green jackfruit flour reduces WDSW-induced inflammation and fibrosis and prevents MASH-induced hepatocarcinogenesis.

**Figure 6:**
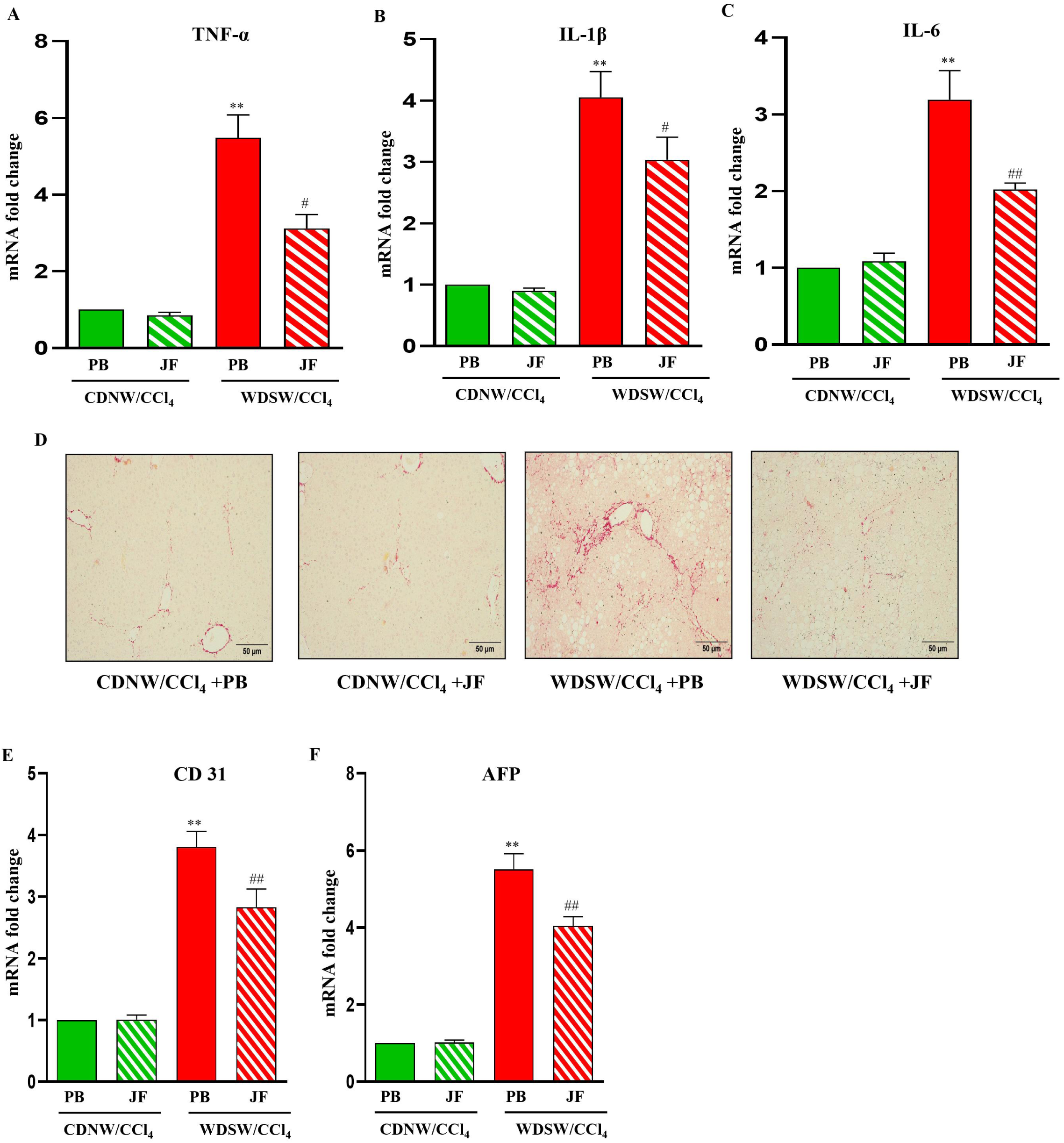
Green jackfruit flour reduces inflammation, fibrosis, and tumorigenesis in the liver of MASH-HCC mice. The liver tissues of mice fed with placebo flour (PB) or green jackfruit flour (JF) from CDNW/CCl_4_ and WDSW/CCl_4_ were processed. The relative hepatic mRNA expression of (A) TNF-α, (B) IL-1β, and (C) IL-6 was determined by quantitative real-time polymerase chain reaction (qRT-PCR). The expression levels were normalized with β-actin. Liver sections were stained for picrosirius red and (D). The relative hepatic mRNA expression of CD31 (E) and AFP (F) was measured by qRT-PCR and normalized with β-actin. All the data are expressed as mean±SEM for 6-8 mice per group. **p<0.001 or ^#^p<0.05 compared to CDNW/CCl_4_ PB; ^##^p<0.001 compared to WDSW/CCl_4_ PB. CDNW, chow diet normal water; WDSW, western diet sugar water; TNF-α, tumor necrosis factor alpha; IL-1β, interleukin 1 beta; IL-6, interleukin 6; CD31, cluster differentiation 31; AFP, alpha-fetoprotein.

### 3.4 Therapeutic effects of green jackfruit flour are mediated via the AMPK and MAPK signaling pathways

In the context of MASLD, dysregulation of the adenosine monophosphate-activated protein kinase (AMPK) pathway has been implicated in the development and progression of the disease [27]. Reduced AMPK activity has been observed in the livers of individuals with MASLD, particularly in those with MASH [28]. This impaired AMPK signaling contributes to metabolic dysregulation, insulin resistance, lipid accumulation, and inflammation in the liver, which are pivotal features of MASLD pathogenesis [29]. Consequently, our study aimed to investigate whether green jackfruit flour could regulate the AMPK pathway, thereby inhibiting lipogenesis and promoting lipid oxidation. To assess this, we examined the hepatic expression of key markers involved in lipid metabolism, such as acetyl-CoA carboxylase (ACC), fatty acid synthase (FASN), and sterol regulatory element binding protein 1 (SREBP1). Interestingly, JF resulted in the downregulation of ACC, FASN, and SREBP1 in WDSW mice, indicating reduced fatty acid synthesis **(Figure 7A-7C)**. Additionally, our findings were corroborated through western blot analysis, revealing increased phosphorylated AMPK (pAMPK) and decreased ACC, FASN, and SREBP1 levels in WDSW+JF mice compared to the placebo group (**Figure 7D)**.

**Figure 7:**
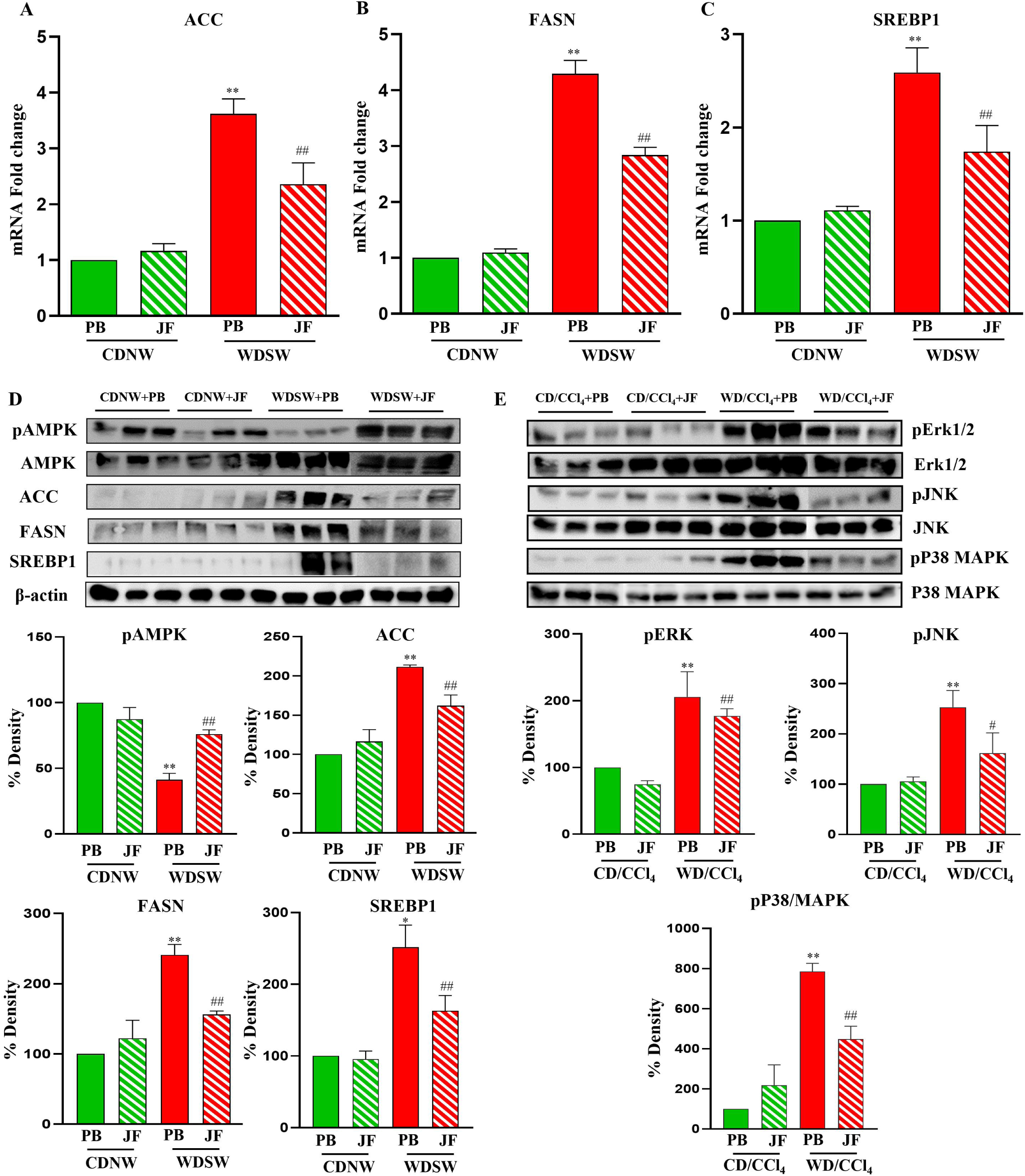
Effect of green jackfruit flour on the AMPK and MAPK signaling pathways. The relative hepatic mRNA expression of (A) ACC, (B) FASN, and (C) SREBP1 was determined by quantitative real-time polymerase chain reaction (qRT-PCR). The expression levels were normalized with β-actin. Western blots were performed on the whole liver tissue lysates of all four groups from the MASH and MASH-HCC experimental models. Representative western blot images and densitometric analysis of (D) pAMPK and AMPK, ACC, FASN, and SREBP1 from the MASH murine model. (E) pErk1/2 and Erk1/2, pJNK and JNK, pP38 MAPK and P38 MAPK from the MASH-HCC murine model. The blots were normalized to the respective endogenous controls. All the data are expressed as mean±SEM for 6-8 mice per group. **p<0.001 compared to CDNW PB or CDNW/CCl_4_ PB; ^##^p<0.001 compared to WDSW PB or WDSW/CCl_4_ PB. CDNW, chow diet normal water; WDSW, western diet sugar water; PB, placebo flour; JF, green jackfruit flour; ACC, acetyl-CoA carboxylase; FASN, fatty acid synthase; SREBP1, sterol regulatory element-binding protein 1; p-ERK1/2, phospho-extracellular signal-regulated protein kinase; p-JNK, phospho-c-Jun N-terminal kinase; pP38 MAPK, phospho-p38 mitogen-activated protein kinase.

In the realm of MASH-associated HCC, dysregulation of the mitogen-activated protein kinase (MAPK) pathway is multifaceted, involving mechanisms such as dysregulation of glucose metabolism and fatty acid synthesis, inflammation, and oxidative stress [30]. Key components of the MAPK pathway, including extracellular signal-regulated kinases (ERK), c-Jun N-terminal kinases (JNK), and p38 MAPK, have been implicated in MASH-HCC pathogenesis [31]. Along the same lines, immunoblot analysis of pErk1/2, pJNK, and pP38 MAPK showed a marked reduction in WDSW+JF mice **(Figure 7E)**. Collectively, our findings provide compelling evidence that green jackfruit flour effectively suppresses hepatic lipogenesis via the activation of the AMPK signaling pathway and prevents the progression to hepatocarcinogenesis by inhibiting MAPK and its downstream effectors.

The therapeutic potential of any diet can indeed be completely based on its nutritional composition. Green jackfruit flour prepared from dehydrated, mature, unripe jackfruits significantly differs from the placebo flour in terms of soluble fibers. Dietary fibers are indigestible portions of plant food that are of prime importance for various beneficial effects. Particularly, soluble fibers serve as a primary source for the production of short-chain fatty acids (SCFAs), products of intestinal microbial anaerobic fermentation [32]. Studies have shown that supplementing SCFAs exogenously can effectively diminish liver fat and enhance metabolic health [33]. This prompted our investigation into the impact of SCFAs from green jackfruit on MASH and HCC. Studies by Koh et al. revealed that fermented jackfruit supplementation in high-fat-fed mice showed an anti-obesity effect through significant improvement in the production of SCFAs, particularly acetic acid and propionic acid [34]. Another study by Zhu et al. showed that the polysaccharide from the jackfruit pulp improved the gut microbiome and significantly increased SCFA production [35]. However, SCFA production specifically attributed to green jackfruit flour remains unexplored. The primary SCFAs produced in the gut are acetate, propionate, and butyrate. Propionate stands out among these, as 80% of propionate in the portal vein is transported to the liver for further metabolism [36]. This led us to hypothesize whether the propionate produced in the gut from the green jackfruit flour is responsible for its therapeutic actions against MASH and HCC.

Initially, we assessed the dose-dependent cytotoxic effects of sodium propionate (NaP) on human hepatic cells (HepG2 cells and Hep3B cells) to choose the optimum concentration for the treatment **(Supplementary Figure 3)**. Subsequently, an *in vitro* model of steatosis was established by treating the human hepatic cells, HepG2, with the free fatty acids sodium palmitate (SP) or sodium oleate (SO). Interestingly, the treatment with sodium propionate on these HepG2 cells resulted in a reduction in intracellular lipid accumulation, as evidenced by oil red o staining **(Figure 8A)**. Furthermore, treatment with NaP upregulated AMPK activation and showed reduced mRNA and protein expression of ACC, FASN, and SREBP1 in the cellular model of MASH **(Figure 8B-8E)**. Consistent with these data, NaP treatment on human HCC cells, Hep3B, and QGY-7703 cells demonstrated reduced expression of pErk1/2, pJNK, and pP38 MAPK **(Figure 8F)**. Thus, green jackfruit flour exerts its therapeutic potential in MASH and MASH-HCC via regulating the AMPK and MAPK signaling pathways, respectively.

**Figure 8:**
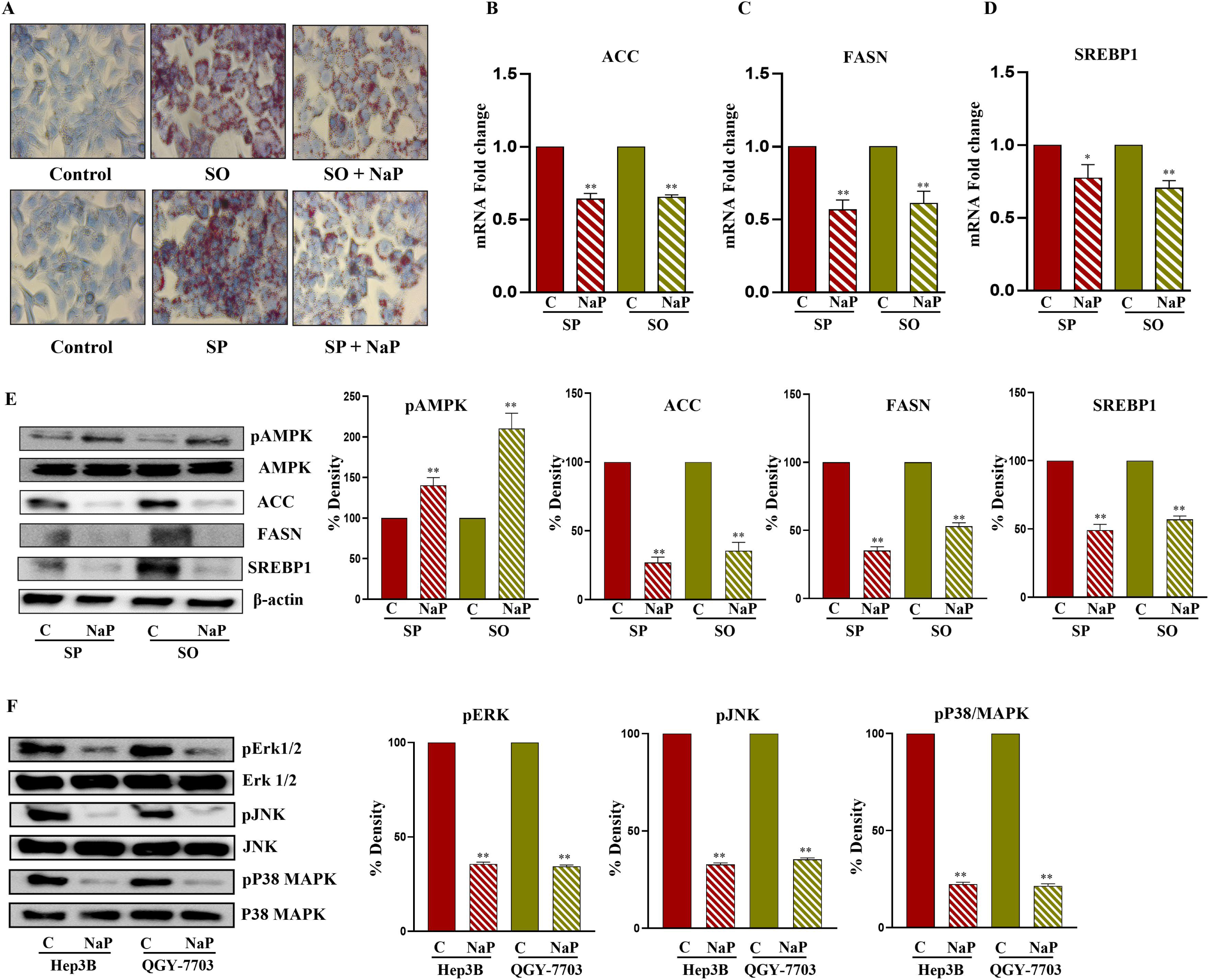
Effect of Sodium propionate treatment on the regulation of AMPK and MAPK signaling pathways in the *in vitro* models of MASH and HCC. HepG2 cells were treated with the free fatty acids sodium palmitate (SP) and sodium oleate (SO). The *in vitro* MASH model was treated with 1 mM of sodium propionate (NaP) for 48 hours. Intracellular lipid accumulation was determined by Oil red O staining (A). The cells were collected, and the relative mRNA expression of (B) ACC, (C) FASN, and (D) SREBP1 was determined by quantitative real-time polymerase chain reaction (qRT-PCR). The expression levels were normalized with β-actin. Western blots were performed on the whole cell lysates of all the groups. Representative western blot images and densitometric analysis of (E) pAMPK and AMPK, ACC, FASN, and SREBP1 from the *in vitro* MASH model. Human HCC cells, Hep3B cells, and QGY-7703 cells were treated with 4 mM NaP for 48 hours and whole cell lysates were prepared for western blotting. Immunoblots for (F) pErk1/2 and Erk1/2, pJNK and JNK, pP38 MAPK, and P38 MAPK. The blots were normalized to the respective endogenous controls. All the data are expressed as mean±SEM. **p<0.001 or *p<0.05 compared to controls. SP, sodium palmitate; SO, sodium oleate; NaP, sodium propionate; ACC, acetyl-CoA carboxylase; FASN, fatty acid synthase; SREBP1, sterol regulatory element-binding protein 1; p-ERK1/2, phospho-extracellular signal-regulated protein kinase; p-JNK, phospho-c-Jun N-terminal kinase; pP38 MAPK, phospho-p38 mitogen-activated protein kinase.

## 4. Discussion

MASLD (formerly known as NAFLD), a non-stigmatizing nomenclature and diagnosis, is a highly prevalent liver disorder affecting more than 30% of the population worldwide [3,4]. MASH represents a more severe form of the disease spectrum and is associated with an increased risk of developing hepatocellular carcinoma (HCC), the most common type of primary liver cancer worldwide [5]. The increased prevalence of MASLD and its progression to MASH and MASH-related HCC pose significant challenges to global public health. Therapeutic options for MASH and MASH-HCC are currently limited, with lifestyle modifications being the cornerstone of management [6]. Diet and exercise play pivotal roles in the treatment of MASH, as they can effectively improve metabolic parameters, promote weight loss, and reduce hepatic fat accumulation [8]. Low-carbohydrate diets, mediterranean diets, and plant-based diets have all shown promise in improving liver function and histology among the numerous dietary regimens studied for the treatment of MASH [9]. Low-carbohydrate diets, characterized by reduced intake of carbohydrates and increased consumption of protein and healthy fats, have been shown to decrease hepatic fat content and improve insulin sensitivity in individuals with MASH [37]. Similarly, mediterranean diets rich in fruits, vegetables, whole grains, and healthy fats like olive oil and nuts have been shown to improve liver enzymes and histology in individuals with MASLD [38]. Plant-based diets, which emphasize fruits, vegetables, legumes, and whole grains while reducing animal products, have also been linked to decreased hepatic fat deposition and enhanced liver function [39].

Previous studies have highlighted the therapeutic potential of jackfruit (*Artocarpus heterophyllus*) and its bioactive compounds for various health conditions [13]. However, it is worth noting that the majority of this research has focused on specific bioactive constituents rather than assessing the therapeutic efficacy of jackfruit flour as a comprehensive dietary supplement [40,41]. Jackfruit is a tropical vegetable and fruit known for its rich nutritional profile and bioactive constituents, including phytochemicals, antioxidants, and dietary fiber [42]. Interestingly, clinical investigations employing the use of green jackfruit flour have unveiled its potential in ameliorating HbA1c levels, reducing plasma glucose levels, and enhancing glycemic control. This effect is attributed to its high fiber content and low net carbohydrate composition, distinguishing it from the placebo flour [20]. Furthermore, a separate study utilizing the same green jackfruit flour explored its impact on chemotherapy-induced leukopenia (CIL), revealing that dietary supplementation with green jackfruit effectively prevented CIL in individuals receiving pegfilgrastim. This protective mechanism is ascribed to the presence of pectin, a natural antioxidant found in green jackfruit, which aids in tumor prevention [21]. Despite the growing body of evidence supporting the therapeutic potential of jackfruit and its derivatives, further research is warranted to elucidate the mechanisms of action and evaluate their efficacy as dietary supplements for human health. As a first step towards this goal, in this study, we investigated the potential therapeutic effects of green jackfruit flour on preventing MASH and progression to HCC in experimental models.

The study’s significance is underscored by its utilization of two distinct preclinical murine models that closely mimic the spectrum of human disease in terms of histology, metabolic parameters, and transcriptomic signatures, rendering them ideal murine models for the study as opposed to several models that may not accurately replicate the disease pattern. Specifically, we employed a diet-induced steatosis and steatohepatitis model, as well as a diet supplemented with chemical induction of metabolic dysfunction-associated steatohepatitis (MASH)-associated hepatocellular carcinoma (HCC), each spanning a study period of 24 weeks [43,44]. In order to ensure the integrity and reliability of this preclinical study, the green jackfruit flour and placebo flour (wheat flour) were purchased from authentic sources manufactured by companies that follow globally benchmarked stringent quality control standards set by FSSAI (Food Safety and Standards Authority of India). In the current study, we employed the use of 5 kcal% of green jackfruit with an equal volume of placebo flour in the chow diet and western diet, which were formulated and customized by Research Diets, Inc., USA.

To the best of our knowledge, this is the first preclinical study that demonstrated that JF effectively reduced obesity, liver injury and insulin resistance in MASH and prevented tumorigenesis in MASH-HCC. This was evidenced by the significant reduction in hepatic steatosis, inflammation, fibrosis, and tumor markers. Notably, WDSW+JF mice showed a significant decrease in the expression of pro-inflammatory cytokines, indicating its potent anti-inflammatory properties. Thus, these data suggest that a 5% calorie replacement with green jackfruit flour could prevent obesity, neutralizing the obesogenic properties of its added sugar and unhealthy fat. These observations align with previous studies suggesting the anti-inflammatory and anti-tumor potential of bioactive compounds found in jackfruit [15,17]. Furthermore, our study elucidates the underlying molecular mechanisms through which JF exerts its hepatoprotective effects. We observed the activation of the AMPK signaling pathway in WDSW+JF mice when compared to WDSW+placebo mice. AMPK, known as a master regulator of energy metabolism, plays a crucial role in maintaining cellular homeostasis and energy balance [27]. Activation of AMPK has been associated with improved lipid metabolism, enhanced insulin sensitivity, and reduced inflammation in the liver [29]. Our findings suggest that the activation of AMPK by JF contributes to its beneficial effects in MASH by regulating lipid metabolism and suppressing inflammatory responses. Moreover, our data also indicate that JF modulates the MAPK signaling pathway, which is implicated in cell proliferation, apoptosis, and inflammation [31]. Specifically, we observed a suppression of MAPK signaling resulting in the reduction of phosphorylation of the downstream effectors, such as ERK1/2, JNK, and p38. Thus, the green jackfruit flour, in addition to reducing hepatic inflammation and fibrosis, also impedes the development of tumorigenesis.

Of note, green jackfruit flour differs significantly from the placebo in its dietary fiber composition [20] and it is also evident that unripe jackfruit has a higher fiber content when compared to ripe jackfruit [45]. Dietary fibers, encompassing a variety of plant-derived carbohydrates resistant to digestion by human enzymes, play a pivotal role in MASLD management [46]. These fibers are known for their ability to modulate gut microbiota composition, promote satiety, and regulate glucose and lipid metabolism [47]. Of particular interest are soluble fibers, such as pectin found abundantly in green jackfruit flour, which undergo fermentation in the colon to produce short-chain fatty acids (SCFAs) [32]. SCFAs, primarily acetate, propionate, and butyrate, are the end products of the microbial fermentation of dietary fibers in the colon. These SCFAs not only serve as an important energy source for colonic epithelial cells but also exert systemic effects on various metabolic processes [48]. In the context of MASLD, SCFAs have been shown to confer numerous benefits, including modulation of hepatic lipid metabolism, attenuation of inflammation, and improvement of insulin sensitivity [33]. Moreover, SCFAs, particularly butyrate, have been implicated in maintaining gut barrier integrity and reducing gut permeability, thereby mitigating the influx of microbial-derived toxins into the systemic circulation [49]. While the majority of butyrate is metabolized in the colon, approximately 40% of acetate and 80% of propionate enter the liver via the portal vein [36]. Additionally, acetate and propionate have been shown to modulate hepatic lipid metabolism and improve insulin sensitivity, further underscoring the importance of SCFAs in MASLD management [50,51]. Intriguingly, previous studies have shown that jackfruit supplementation elevates the levels of SCFAs, in particular propionate, as demonstrated by the caecal GC/MS analysis [34,35]. While the current study did not directly measure the levels of these short-chain fatty acids (SCFAs) in the mice cecum or serum, our future directions aim to expand upon these findings. Our broader plan includes conducting a clinical study and implementing a multi-omics approach to investigate the metabolomic and metagenomic alterations induced by the dietary intervention of green jackfruit flour.

Propionate, the majorly absorbed SCFA that is metabolized in the liver, plays a significant role in modulating the metabolic processes of the liver [51,52]. Propionate has been shown to attenuate hepatic lipogenesis, or the production of fats in the liver, by inhibiting key enzymes involved in this process [51]. Propionate has been linked to improvements in glucose metabolism and insulin sensitivity, which are crucial for maintaining liver health [51,53]. Furthermore, emerging evidence suggests that propionate may also exert anti-inflammatory and antiapoptotic effects [54]. While these pathways indicate that propionate may play a role in lowering the risk of HCC, further research is needed to completely understand its effects and establish the best strategies for employing propionate or its precursors (sources rich in soluble dietary fiber) to prevent HCC. Green jackfruit may continue to garner translational attention as researchers investigate how it modulates the gut microbiota and the role of propionate in promoting gut barrier integrity, maintaining a healthy gut-liver axis, and potentially reducing HCC development. Our data illustrates the impact of propionate on an *in vitro* model of MASH and HCC. Consistent with our *in vivo* findings, external treatment with sodium propionate markedly regulated the AMPK pathway in the MASH model and the MAPK pathway in the HCC model. Thus, the current investigation also sets the stage for confirmation in clinical studies.

In conclusion, our findings provide compelling evidence for the therapeutic potential of JF in improving metabolic health, preventing MASH and preventing its progression to HCC via the AMPK and MAPK signaling pathways. Furthermore, our findings highlight the importance of short-chain fatty acids, particularly propionate, in modulating liver metabolism and inflammation, emphasizing the potential of dietary interventions that modulate gut microbiota-derived metabolites, which play an important role in MASLD. Further clinical studies are in the pipeline employing multi-omics approaches to elucidate the comprehensive metabolic and molecular alterations induced by green jackfruit intervention, which would pave the way for the development of innovative dietary strategies for the prevention and treatment of MASLD.

**Figure 9:**
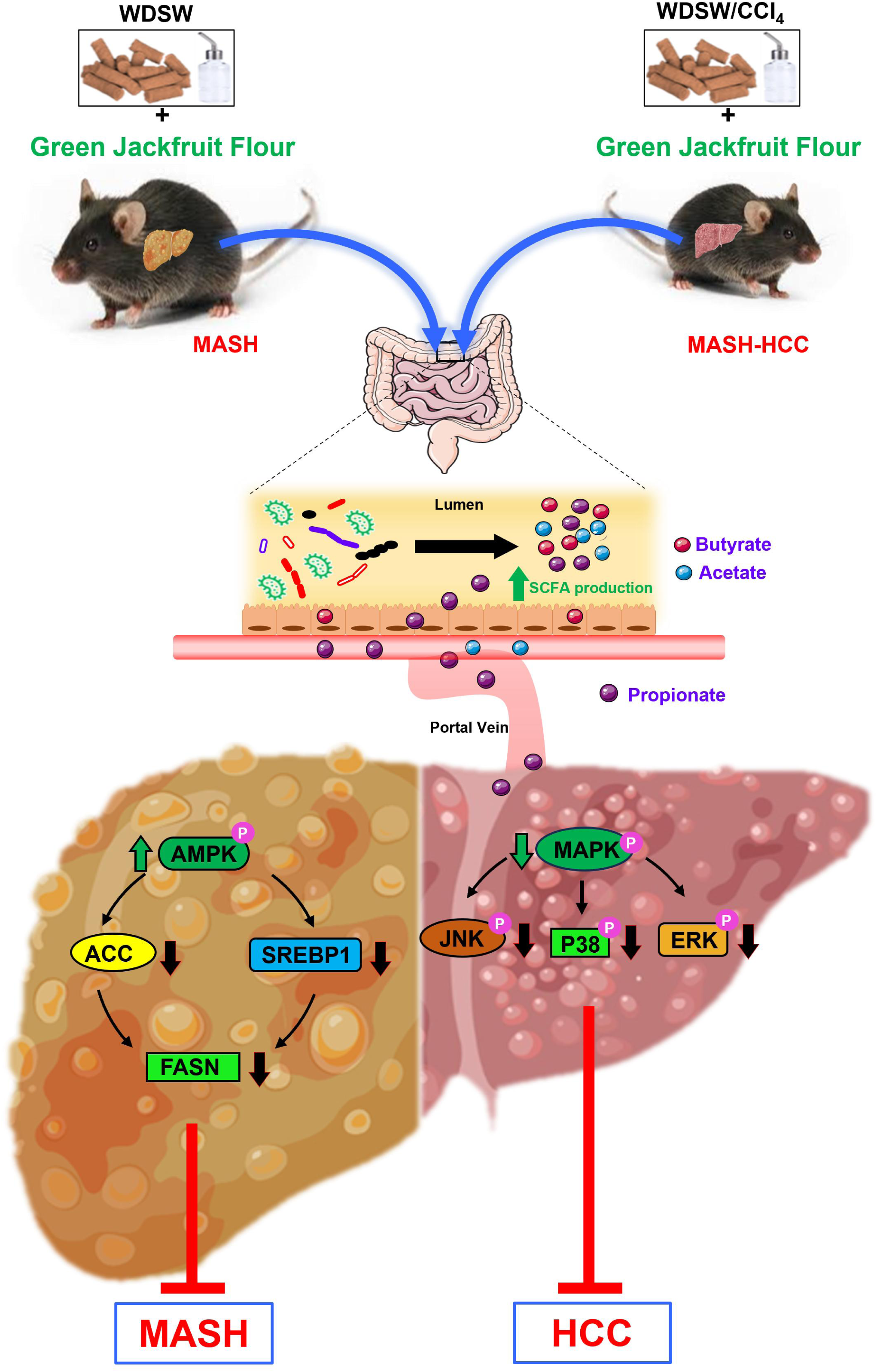
Molecular mechanisms involved in the therapeutic action of green jackfruit flour in the prevention of MASH and progression to HCC in experimental models. MASH and MASH-HCC mice were treated with 5 kcal% of green jackfruit flour for 12 weeks. The green jackfruit flour, rich in soluble fibers, undergoes fermentation in the colon to produce short-chain fatty acids (SCFA-acetate, propionate, and butyrate). The majority of butyrate is metabolized in the colon, whereas acetate (40%) and propionate (80%) enter the liver via the portal vein. Propionate, via activation of AMPK and inhibition of MAPK, prevents metabolic dysfunction-associated steatohepatitis and progression to hepatocarcinogenesis, respectively.

## CRediT authorship contribution statement

**Bharathwaaj, Diwakar, and Akshatha:** Investigation, Data curation and analysis, Writing-original draft, Writing-review and editing. **Amith Bharadwaj:** Methodology, Investigation. **James Joseph:** Conceptualization, Writing-review and editing, Funding Acquisition. **Megha:** Conceptualization, Writing-review and editing. **Suchitha Satish:** Investigation. **Deepak Suvarna:** Investigation. **Prasanna K. Santhekadur:** Data analysis, Writing-review and editing. **Saravana B. Chidambaram:** Data analysis, Writing-review and editing. **Ajay Duseja:** Conceptualization, Writing-review and editing. **Divya P. Kumar:** Conceptualization, Investigation, Validation, Supervision, Funding acquisition, Writing-original draft, review and editing.

## Data availability

Data will be made available on request.

## Acknowledgements

This study was supported in whole or in part, by the SERB-POWER grant (SPG/2021/002524) and God’s Own Food Solutions Pvt Ltd, India to DPK, and Senior Research Fellowship (SRF) from ICMR to ANS.

The authors also acknowledge funding support from the Department of Biotechnology-Boost to University Interdisciplinary Life Science Departments for Education and Research programme (DBT-BUILDER: BT/INF/22/SP43045/2021) and the Department of Science and Technology–Promotion of University Research and Scientific Excellence (DST-PURSE) (SR/PURSE/2021/81 (c).

## Abbreviations

MASLD: metabolic dysfunction-associated steatotic liver disease
NAFLD: nonalcoholic Fatty liver disease
MASL: metabolic dysfunction-associated steatotic liver
MASH: metabolic dysfunction-associated steatohepatitis
NASH: non-alcoholic steatohepatitis
HCC: hepatocellular carcinoma
JF: jackfruit flour
PB: placebo flour
CDNW: chow diet normal water
WDSW: western diet sugar water
FDA: food and drug administration
CIL: chemotherapy-induced leukopenia
AST: aspartate transaminase
ALT: alanine transaminase
TG: total triglycerides
GTT: glucose tolerance test
ITT: insulin tolerance test
HOMA-IR: homeostatic model assessment of insulin resistance
DMEM: dulbecco’s modified eagle medium
SP: Sodium Palmitate
SO: Sodium Oleate
NaP: sodium propionate
AMPK: adenosine monophosphate-activated protein kinase
MAPK: mitogen-activated protein kinase
JNK: c-Jun N-terminal kinases
ERK1/2: extracellular signal-regulated kinase 1/2
ACC: Acetyl-CoA carboxylase
FASN: Fatty acid synthase
SREBP1: Sterol Regulatory Element-binding Protein-1
P38 MAPK: p38 Mitogen-activated protein kinase
ECL: enhanced chemiluminescence
HRP: horseradish peroxidase
TNF-α: tumor necrosis factor-alpha
IL-6: interleukin-6
IL-1β: interleukin-1β
CD31: cluster differentiation 31
AFP: alpha-fetoprotein
SCFA: short-chain fatty acids.

---

**Supplementary Figure 1:**
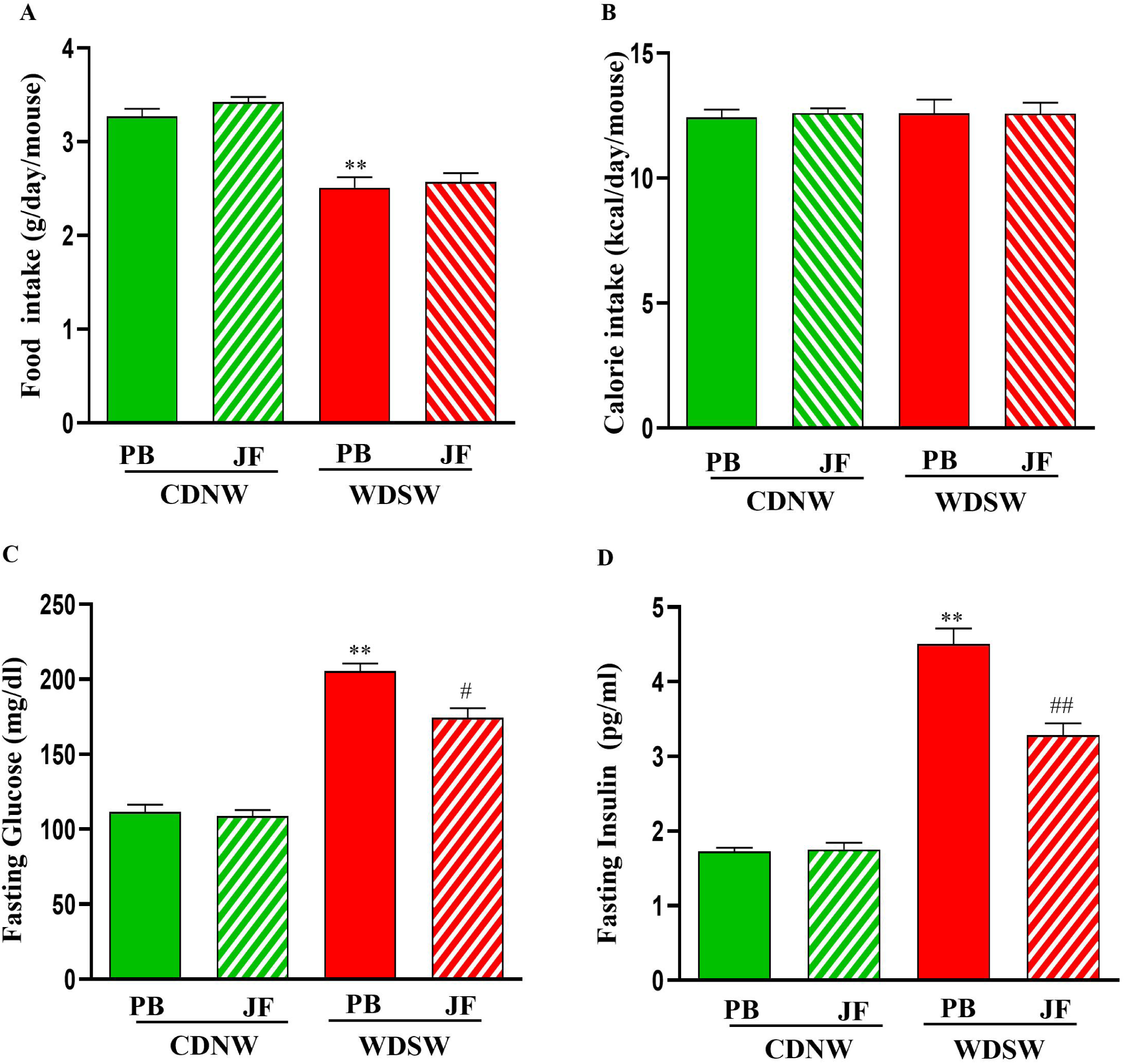
Effect of green jackfruit flour on food intake, calorie intake, fasting glucose, and fasting insulin in experimental MASH model. Mice were fed with either CDNW or WDSW for 12 weeks. Further, mice were randomized into four groups based on dietary interventions. Mice were fed with CDNW with placebo flour (PB) or green jackfruit flour (JF) and WDSW with placebo flour or green jackfruit flour for an additional 12 weeks. Food intake (A), calorie intake (B), fasting glucose (C), and fasting insulin (D) were measured. All the data are expressed as mean±SEM for 6-8 mice per group. **p<0.001 compared to CDNW PB; ^##^p<0.001 compared to WDSW PB.

**Supplementary Figure 2:**
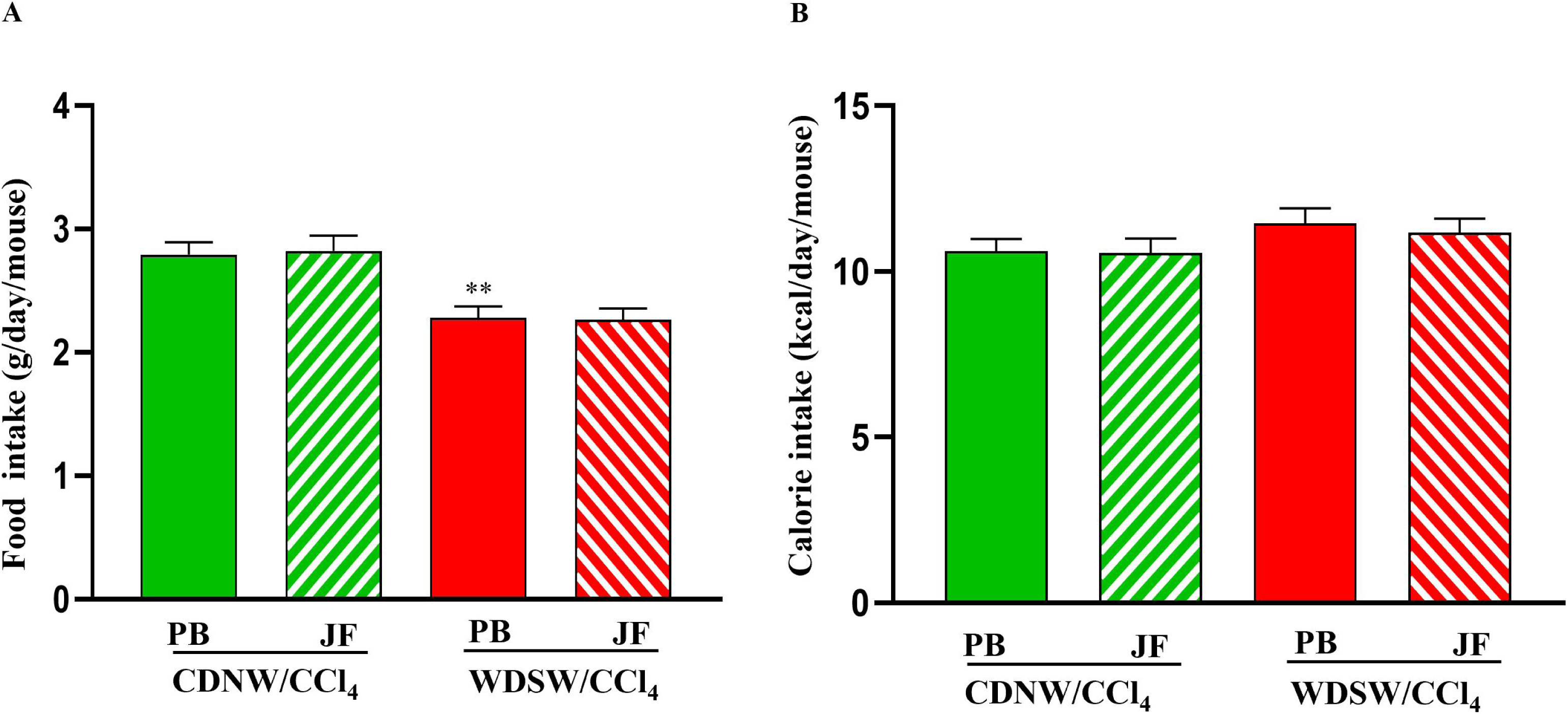
Monitoring of food intake and calorie intake upon green jackfruit flour dietary intervention in MASH-HCC murine model. CDNW/CCl_4_ and WDSW/CCl_4_ mice were fed with placebo flour (PB) or green jackfruit flour (JF) for 12 weeks. Food intake (A) and calorie intake (B) were monitored. All the data are expressed as mean±SEM for 6-8 mice per group. **p<0.001 compared to CDNW PB; ^##^p<0.001 compared to WDSW PB.

**Supplementary Figure 3:**
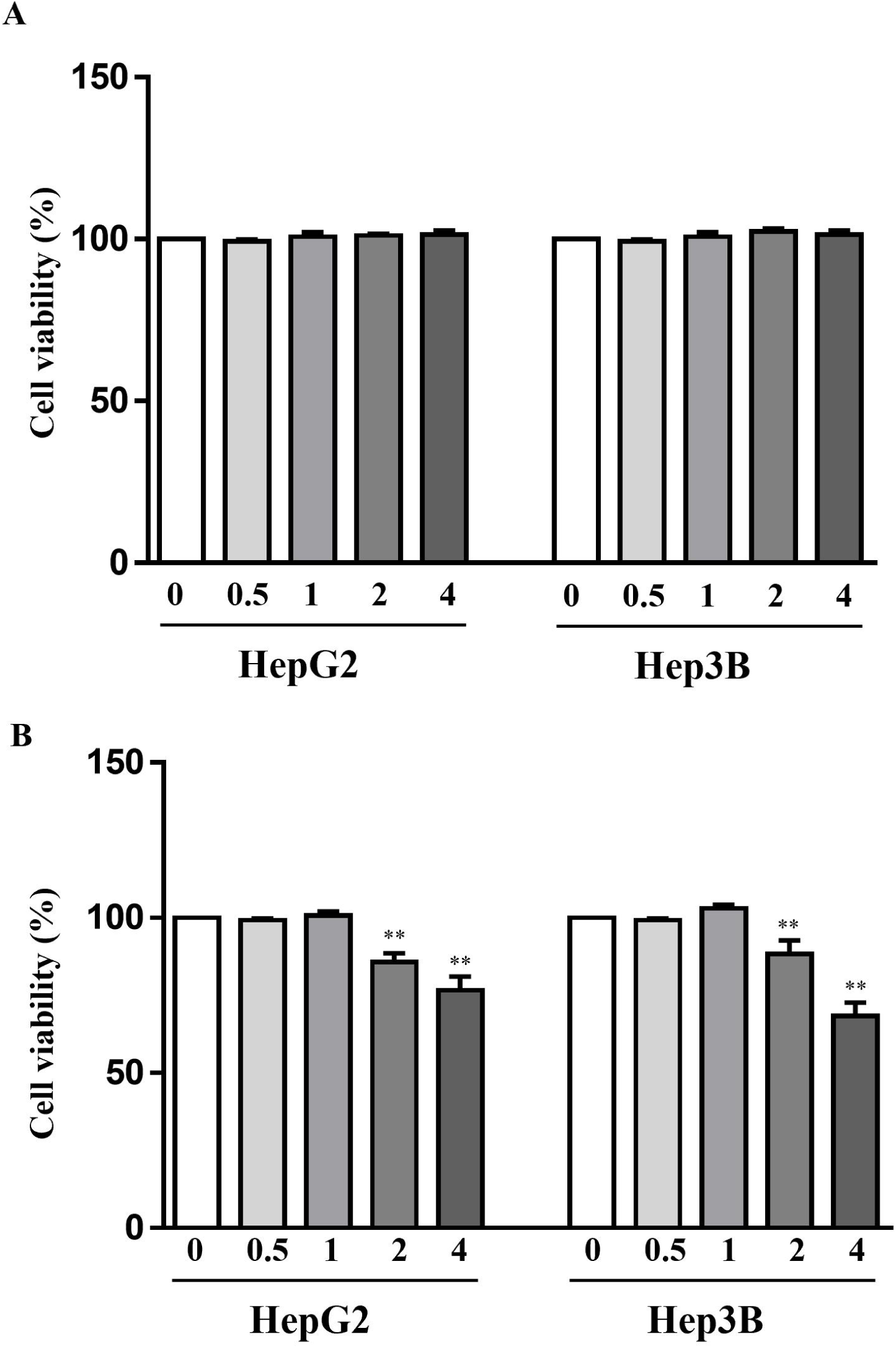
Effect of sodium propionate on cell viability of HepG2 and Hep3B cells. WST-1 assay was performed to assess the cell viability of sodium propionate (NaP)-treated groups compared to the control groups. HepG2 and Hep3B cells were treated with 0, 0.5, 1, 2, or 4mM NaP for (A) 24 hrs and (B) 48 hrs. All the data are expressed as mean±SEM. **p<0.001 compared to control.

**Supplementary Table 1:**
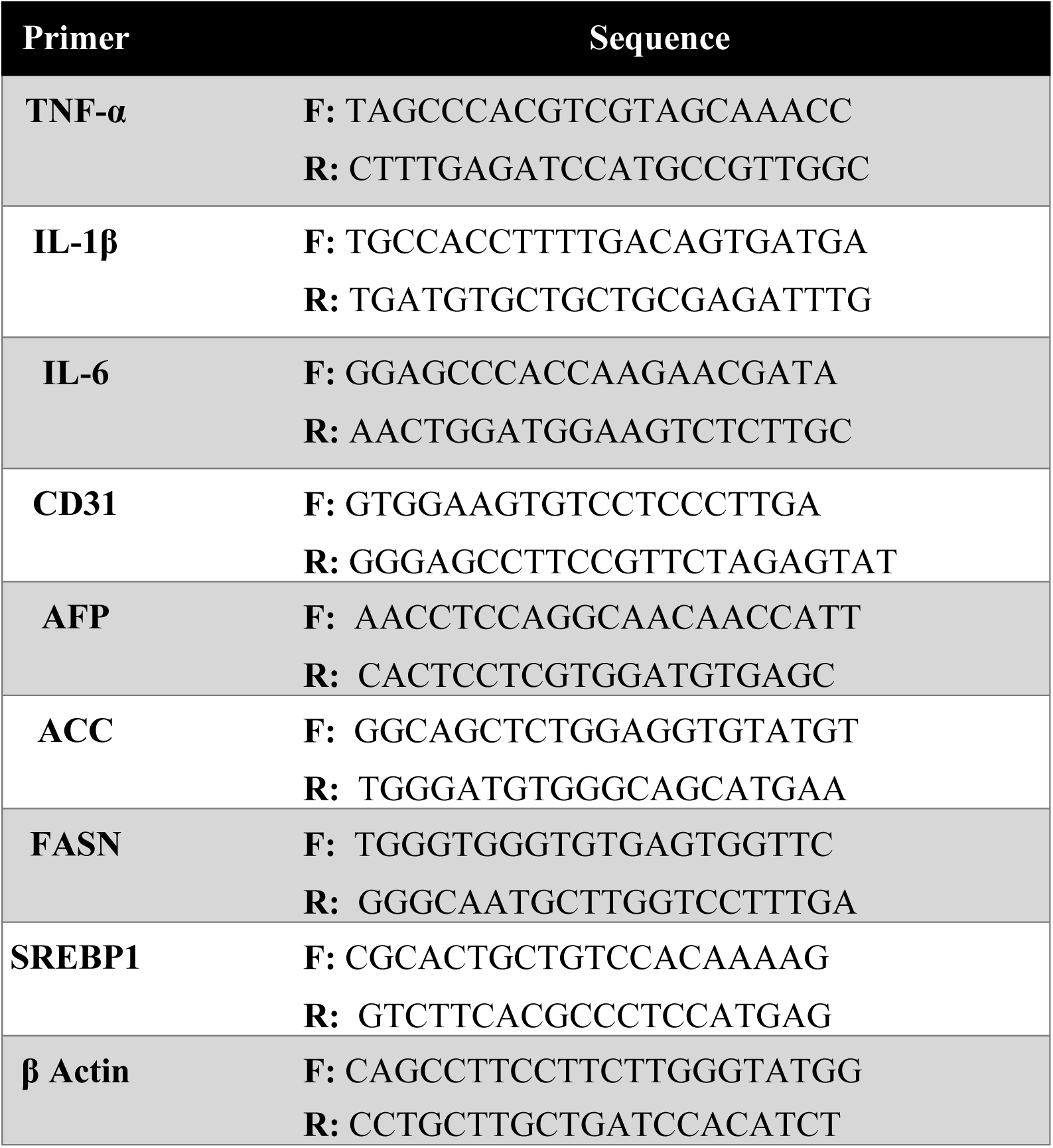
List of primer sequences used in qRT-PCR.

